## Supplementary material for "Causal role of the dorsolateral prefrontal cortex in modulating the balance between Pavlovian and instrumental systems in the punishment domain": S1 and S2 Tables

**S1 Table**. Side effects

|  | Sham  (# session = 42) | Anode  (# session = 51) | Total  (# session = 93) | p-value |
| --- | --- | --- | --- | --- |
| tingling | 0.714 (0.944) | 1.176 (1.126) | 0.968 (1.068) | 0.037 |
| itching | 0.738 (1.037) | 0.824 (1.178) | 0.785 (1.112) | 0.714 |
| skin irritation | 0.690 (1.137) | 0.490 (0.946) | 0.581 (1.035) | 0.356 |
| skin pain | 0.405 (0.665) | 0.373 (0.747) | 0.387 (0.708) | 0.828 |
| headache | 0.310 (0.780) | 0.000 (0.000) | 0.140 (0.544) | 0.006 |
| fatigue | 1.333 (1.391) | 1.098 (1.063) | 1.204 (1.221) | 0.358 |
| difficulty  concentration | 1.405 (1.466) | 0.843 (1.046) | 1.097 (1.277) | 0.034 |
| mood  disturbance | 0.119 (0.328) | 0.196 (0.601) | 0.161 (0.495) | 0.459 |
| visual  distortion | 0.333 (0.786) | 0.157 (0.505) | 0.237 (0.649) | 0.194 |

* scale 0 - 5

**S2 Table**. Manipulation check of double-blind design

|  | **Sham**  **(# session = 42)** | **Anode**  **(# session = 51)** | **Total**  **(# session = 93)** | **p-value** |
| --- | --- | --- | --- | --- |
| continuity |  |  |  | < 0.001 |
| continue | 4 (9.5%) | 22 (43.1%) | 26 (28.0%) |  |
| discontinue | 38 (90.5%) | 29 (56.9%) | 67 (72.0%) |  |
| duration |  |  |  | < 0.001 |
| less than 5 mins | 30 (79.0%) | 16 (55.2%) | 46 (68.7%) |  |
| less than 10 mins | 6 (15.8%) | 8 (27.6%) | 14 (20.9%) |  |
| less than 15 mins | 2 (5.3%) | 5 (17.2%) | 7 (10.5%) |  |
| direction |  |  |  | 0.47 |
| toward cheek | 16 (38.1%) | 25 (49.0%) | 41 (44.1%) |  |
| far from cheek | 8 (19.0%) | 6 (11.8%) | 14 (15.1%) |  |
| don't know | 18 (42.9%) | 20 (39.2%) | 38 (40.9%) |  |

* Continuity: did you feel that the electric current continued to flow during the tDCS experiment?

* Duration: if not, how long did you feel that the electric current flowed?

* Direction: did you feel that the current flowed toward your cheek or far from your cheek?
