## Supplementary material for "Causal role of the dorsolateral prefrontal cortex in modulating the balance between Pavlovian and instrumental systems in the punishment domain": S1 file

**Electric shock**

The electric shock was tailored to each participant at a level of ‘moderately unpleasant’ for safety. Each participant conducted a task consisting of 3 blocks to find the level. In the first block, electric stimulation of direct currents below 7 mA was first applied, and participants rated their subjective feelings on an 11-point scale (0 = not unpleasant at all, 10 = very unpleasant). If the participants rated it as 6-10 points, the intensity of the next stimulation decreased, otherwise increased. The amount of change in stimulation was halved each time it changed. When the amount of change was less than 0.1 mA, the first block ended [[1–5]](https://paperpile.com/c/CBwJ4k/dgYP+dYZM+niTD+HeUn+Gvks). The second and third blocks had the same process as the first, and only the currents initially given were less than 6 mA and less than 8 mA, respectively. By substituting a series of electrical stimulation intensities into the sigmoid function, the intensity corresponding to the subjective intensity of 5 points (moderately unpleasant) was estimated. No matter what, the current higher than 12.4 mA was not used throughout the process [[6]](https://paperpile.com/c/CBwJ4k/Zm7T). The participants were fully informed that they can stop the process at any time (i.e. discomfort during the electrical stimulation experiment).
