## Supplementary figures and images for "Causal role of the dorsolateral prefrontal cortex in modulating the balance between Pavlovian and instrumental systems in the punishment domain"

### S1 Fig

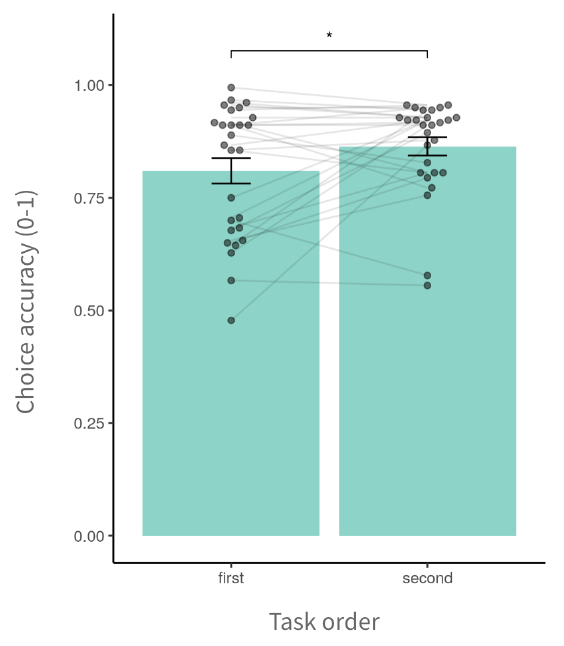

### S2 Fig

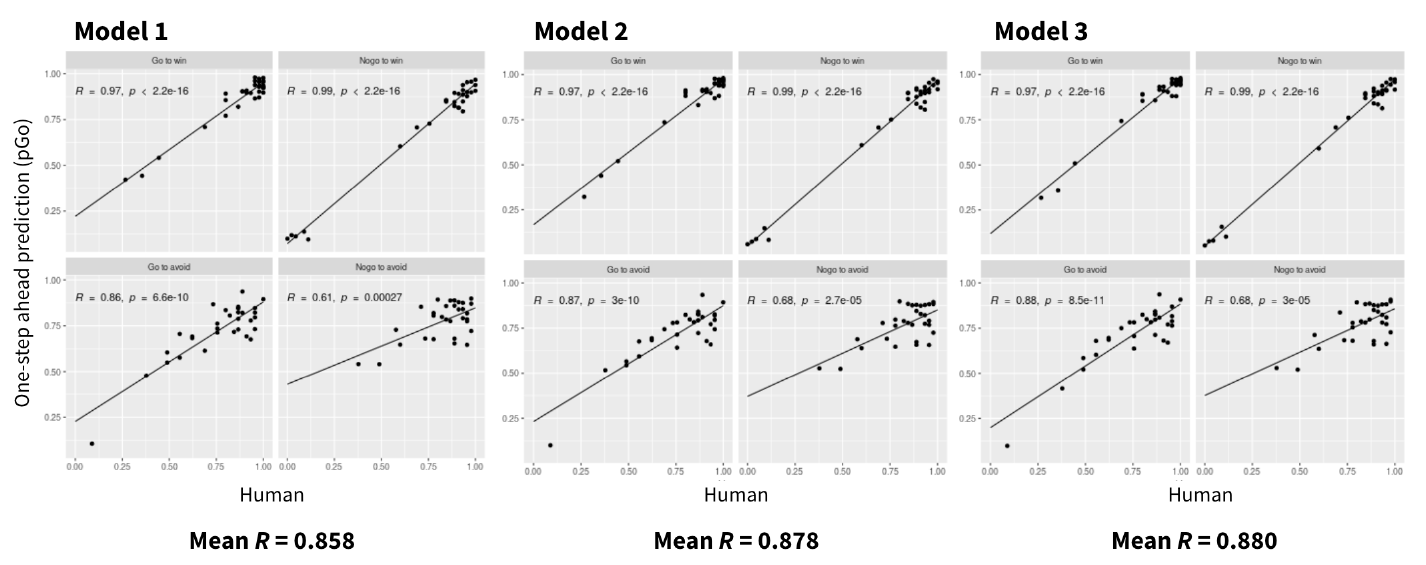

### S3 Fig

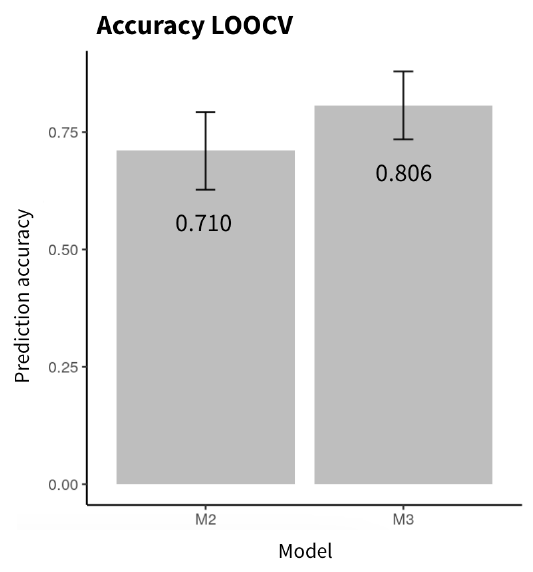
